# Horizontal gene transfer and recombination analysis of SARS-CoV-2 genes helps discover its close relatives and shed light on its origin

**DOI:** 10.1101/2020.12.03.410233

**Authors:** Vladimir Makarenkov, Bogdan Mazoure, Guillaume Rabusseau, Pierre Legendre

## Abstract

**Background:** The SARS-CoV-2 pandemic is among the most dangerous infectious diseases that have emerged in recent history. Human CoV strains discovered during previous SARS outbreaks have been hypothesized to pass from bats to humans using intermediate hosts, e.g. civets for SARS-CoV and camels for MERS-CoV. The discovery of an intermediate host of SARS-CoV-2 and the identification of specific mechanism of its emergence in humans are topics of primary evolutionary importance. In this study we investigate the evolutionary patterns of 11 main genes of SARS-CoV-2. Previous studies suggested that the genome of SARS-CoV-2 is highly similar to the horseshoe bat coronavirus RaTG13 for most of the genes and to some Malayan pangolin coronavirus (CoV) strains for the receptor binding (RB) domain of the spike protein.

**Results:** We provide a detailed list of statistically significant horizontal gene transfer and recombination events (both intergenic and intragenic) inferred for each of 11 main genes of the SARS-Cov-2 genome. Our analysis reveals that two continuous regions of genes S and N of SARS-CoV-2 may result from intragenic recombination between RaTG13 and Guangdong (GD) Pangolin CoVs. Statistically significant gene transfer-recombination events between RaTG13 and GD Pangolin CoV have been identified in region [1215-1425] of gene S and region [534-727] of gene N. Moreover, some significant recombination events between the ancestors of SARS-CoV-2, RaTG13, GD Pangolin CoV and bat CoV ZC45-ZXC21 coronaviruses have been identified in genes ORF1ab, S, ORF3a, ORF7a, ORF8 and N. Furthermore, topology-based clustering of gene trees inferred for 25 CoV organisms revealed a three-way evolution of coronavirus genes, with gene phylogenies of ORF1ab, S and N forming the first cluster, gene phylogenies of ORF3a, E, M, ORF6, ORF7a, ORF7b and ORF8 forming the second cluster, and phylogeny of gene ORF10 forming the third cluster.

**Conclusions:** The results of our horizontal gene transfer and recombination analysis suggest that SARS-Cov-2 could not only be a chimera resulting from recombination of the bat RaTG13 and Guangdong pangolin coronaviruses but also a close relative of the bat CoV ZC45 and ZXC21 strains. They also indicate that a GD pangolin may be an intermediate host of SARS-CoV-2.

## Background

The recent outbreak of a serious pneumonia disease caused by the SARS-CoV-2 (i.e. COVID-19) pathogen has highlighted the danger of coronavirus spread between different zoonotic sources. Some important transfers of genetic information across species have been observed during the first SARS outbreak, involving species from various wet markets in China (Xu et al. 2004). Several recent studies have suggested that the only close relative of SARS-CoV-2 is the RaTG13 CoV found in *Rhinolophus affinis* (horseshoe bats) (Guo et al. 2020; Zhou et al. 2020). Thus, these bats could be considered as the main natural reservoir of the SARS-CoV and SARS-CoV-2 viruses. However, recent analyses of the SARS-CoV-2 genome conducted by Lu et al. (2020) and Lam et al. (2020) have indicated its high resemblance in certain regions with different coronavirus genomes of Malayan pangolins (*Manis javanica*). Zhang et al. (2020) have reported that the SARS-CoV-2 genome is 91.02% identical to that of a Guangdong (GD) Pangolin CoV (virus found in dead Malayan pangolins in the Guangdong province of China; Liu et al. 2019). While at the whole-genome level RaTG13 remains the closest to SARS-CoV-2 coronavirus organism overall (these CoVs share 96% of whole genome identity), the receptor binding (RB) domain of the spike (S) protein of SARS-CoV-2 is much more similar to the RB domain of the GD Pangolin CoV than to that of RaTG13 (Lam et al. 2020; Zhang et al. 2020). Five key amino acid residues taking part in the interaction with human angiotensin-converting enzyme 2 (ACE2) are completely identical in SARS-CoV-2 and GD Pangolin CoV. However, these amino acids are different in the RB domain of RaTG13 and the Guangxi (GX) Pangolin CoV (virus found in Malayan pangolins in the Guangxi province of China). Three possible evolutionary hypotheses could be advanced to explain this paradigm. According to the first one, these mutations may have occurred as a consequence of the phenomenon of parallel evolution, when distinct CoV organisms have undergone similar mutations, and thus have developed similar traits, in response to common evolutionary pressure. For example, it is possible that SARS-CoV-2 had acquired the RB domain mutations during adaptation to passage in cell culture, as has been observed for SARS-CoV (Andersen et al. 2020). The second reasonable hypothesis is divergent evolution favoring amino acid substitutions in the RaTG13 lineage, independent of recombination. Thus, GD Pangolin CoV and SARS-CoV-2 similarity could be the consequence of shared ancestry. According to the third hypothesis, two or more close relatives of SARS-CoV-2 may have been affected by some recombination events within a host species. Such a recombination could result in gene exchange between the CoV genomes. During this recombination, some whole genes of the donor CoV genome could be incorporated into the recipient CoV genome either directly (when the orthologous genes are absent in the recipient) or by supplanting in it the existing orthologous genes. This constitutes the *complete gene transfer model* that accounts for the phenomenon of intergenic recombination (Boc et al. 2010). Moreover, the recombination process could lead to the formation of *mosaic* genes through *intragenic recombination* of the orthologous genes of the donor and recipient CoVs. The term mosaic comes from the pattern of interspersed blocks of sequences with different evolutionary histories. This constitutes the *partial gene transfer model* that accounts for the phenomenon of intragenic recombination (Boc and Makarenkov 2011). These models of reticulate evolution have been widely studied in the literature since the beginning of this century (Legendre 2000; Makarenkov and Legendre 2000; Koonin et al. 2001; Legendre and Makarenkov 2002; Mirkin et al. 2003; Makarenkov and Legendre 2004; Makarenkov et al. 2004; Bruen et al. 2006; Jin et al. 2006; Huson and Bryant 2006; Jin et al. 2007; Glazko et al. 2007; Huson et al. 2010; Glazko et al. 2007; Arenas 2013; Bapteste et al. 2013; Corel et al. 2016).

In this paper, we provide arguments supporting the third hypothesis of the SARS-CoV-2 origin, according to which the SARS-CoV-2 genome is a chimera of the RaTG13 and GD Pangolin coronaviruses. Such a conclusion is in agreement with the recent results of Xiao et al. (2020) and Li et al. (2020). Some authors, however, present evidence that there was not recombination between ancestors of GD Pangolin CoV and RaTG13, suggesting that the similarity pattern between their genomes is rather the result of recombination into RaTG13 from some unknown CoV strains (Boni et al. 2020).

In their study investigating the origins of SARS, Stavrinides and Guttman (2004) highlighted that the SARS-CoV genome is a mosaic of some mammalian and avian virus genomes. These authors also pointed out that recombination between the ancestor viruses of SARS may have happened in the host-determining gene S. However, the work of Stavrinides and Guttman mainly addresses the deep evolutionary origins of the entire SARS CoV clade and has been criticized by some expert in the field (Weiss and Navas-Martin 2005). Hu et al. (2017) have more recently conducted evolutionary analysis of 11 bat CoV (i.e. SARSr-CoV) genomes discovered in horseshoe bats in the Yunnan province of China and found that they share high sequence similarity to SARS-CoV in the hypervariable N-terminal domain and the RB domain of gene S, as well as in some regions of genes ORF3 and ORF8. Hu et al. also reported that their recombination analysis provided evidence of frequent recombination events within genes S and ORF8 between these bat CoVs and suggested that the direct progenitor of SARS-CoV may have originated from multiple recombination events between the precursors of different bat CoVs.

Human CoV strains discovered during previous SARS outbreaks have been often hypothesized to pass from bats to humans using intermediate hosts (e.g. civets for SARS-CoV and camels for MERS-CoV) (Graham and Baric 2010), suggesting that SARS-CoV-2 may have also been transmitted to humans this way. It is worth noting, however, that some other studies from scientists in the field, such as Ralph Baric, Zheng-Li Shi and Peter Dazsak, have showed the potential for direct infection of humans from bat strains. They include both serological studies in rural China near the caves where these bat viruses circulate as well as in vitro/cell culture studies indicating the ability of bat isolates to infect human cells directly (Zhengli 2020). Nevertheless, the discovery of such an intermediate host of SARS-CoV-2, if it existed, is key, as it could shed light on the evolution of this dangerous virus.

At the same time, it is also crucial to retrace the evolution of all genes of the SARS-CoV-2 genome, doing it on a gene-by-gene basis. Evolutionary patterns of various genes of SARS-CoV-2 could be quite different as some of them could be affected by specific horizontal gene transfer and recombination events. These events could witness that the SARS-CoV-2 genome is a mosaic genome obtained via recombination of various virus strains. Our findings suggest that the SARS-CoV-2 genome may be in fact formed via recombination of genomes close to the RaTG13 and GD Pangolin CoV genomes, and be a close relative of bat CoV ZC45 and ZXC21.

## Results

In this section, we present a detailed analysis of putative gene transfer and recombination events that were detected in each of 11 genes of SARS-CoV-2 as well as in the RB domain of the spike protein.

### SimPlot similarity analysis

Our SimPlot analysis (Fig. 1a) conducted with 25 CoV genomes (see the Methods section for a detailed data description) shows that the Wuhan SARS-CoV-2 and RaTG13 genomes share 96.14% of whole-genome identity, while the Wuhan SARS-CoV-2 and GD Pangolin genomes are 90.34% identical. The RaTG13 and GD Pangolin CoV genomes are by far the closest ones to the SARS-CoV-2 genome. For example, only 85.43% of whole-genome identity is shared between the Wuhan SARS-CoV-2 and GX Pangolin CoV genomes. Given such a close resemblance between SARS-CoV-2 and RaTG13, bat is a likely reservoir of origin for SARS-CoV-2, as was the case during previous CoV outbreaks.

**Figure 1.**
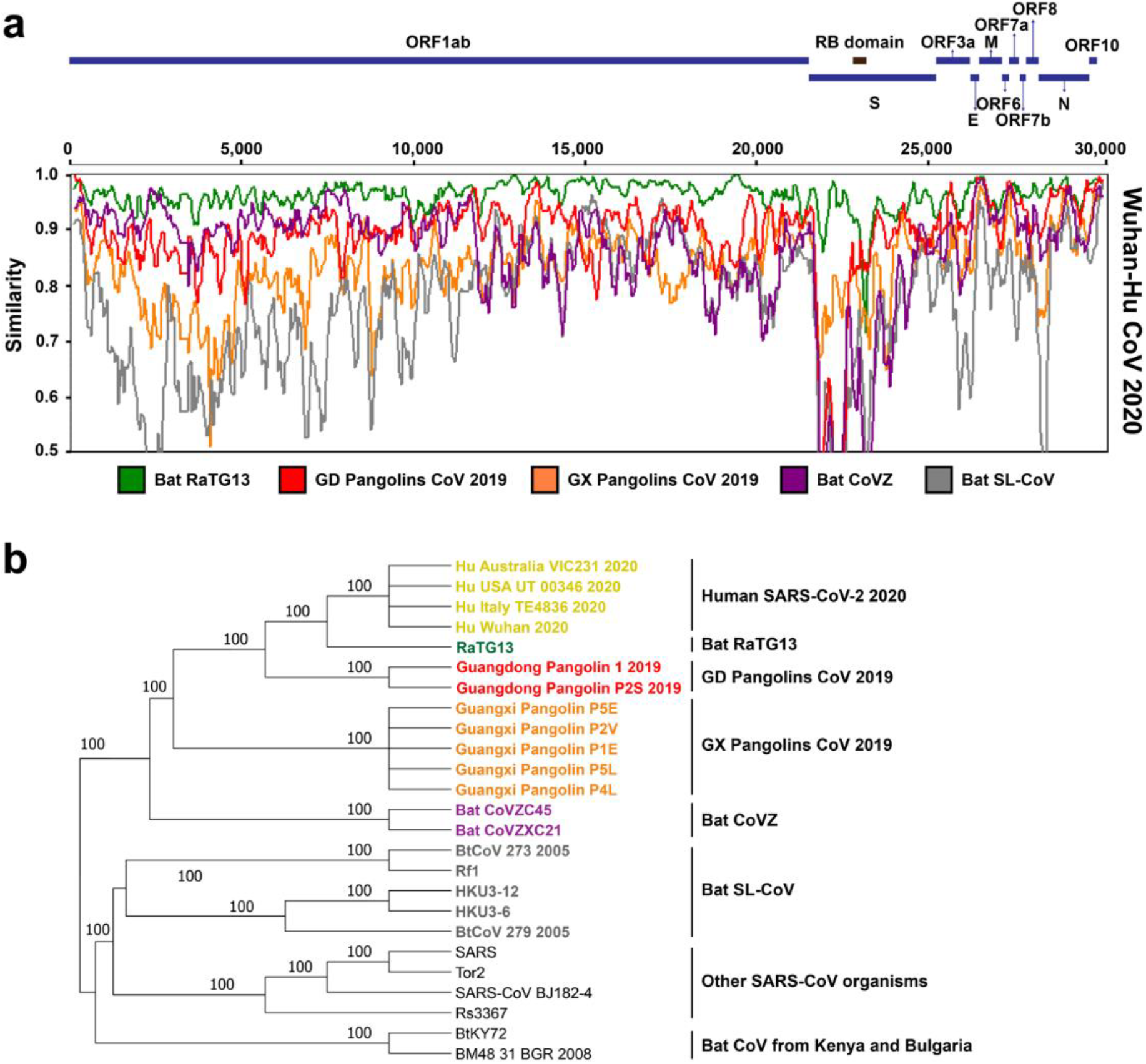
Genome similarity and phylogenetic analysis of SARS-CoV-2 and related viruses: (a) SimPlot sliding window analysis of changing patterns of sequence similarity between: the Wuhan SARS-CoV-2 2020 reference genome with the RatTG13 CoV genome (green) and the consensus genomes of the GD Pangolin CoV (red), GX Pangolin CoV (orange), Bat CoVZ (violet) and Bat SL-VoC (gray) groups. Gene limits for genes ORF1ab, S, ORF3a, E, M, ORF6, ORF7a, ORF7b, ORF8, N and ORF10, as well as for the RB domain, are shown at the top of the figure. Different groups of sequences merged in SimPlot analysis are represented by different colors corresponding to species clusters in the whole genome phylogeny shown in panel (b) of the figure; (b) Whole genome phylogeny of 25 SARS-CoV and SARS-CoV-2-related organisms. Species clusters are indicated on the right. Bootstrap scores are indicated on the internal branches of the tree. Branches with bootstrap score lower than 60% were collapsed. The tree was inferred using the RAxML method with the most suitable for these data, the HKY-gamma evolutionary model, and 100 replicates in bootstrapping.

Then, we performed a detailed similarity analysis at the level of individual genes to compare the Wuhan SARS-CoV-2 gene sequences with the RaTG13, GD Pangolin CoV group sequences and Bat CoVZ group sequences (Fig. 2). Our gene-by-gene analysis revealed some regions where SARS-CoV-2 was more similar to GD Pangolin CoV than to RaTG13. These regions have been found in genes S (at the RB domain level), ORF3a, M, ORF7a and N, suggesting that some recombination events between GD Pangolin CoV and RaTG13 may have occurred not only in gene S (as it has been reported in previous studied; see Lam et al. 2020 and Zhang et al. 2020), but in four other genes of these CoV genomes. We also found that in some continuous gene regions, specifically in genes ORF1ab, ORF3a, M and N, the SARS-CoV-2 gene sequences are much more similar to those of the bat CoV ZC45 and ZXC21 viruses than to those of the GD Pangolin CoVs, and sometimes even of RaTG13 (Fig. 2).

**Figure 2.**
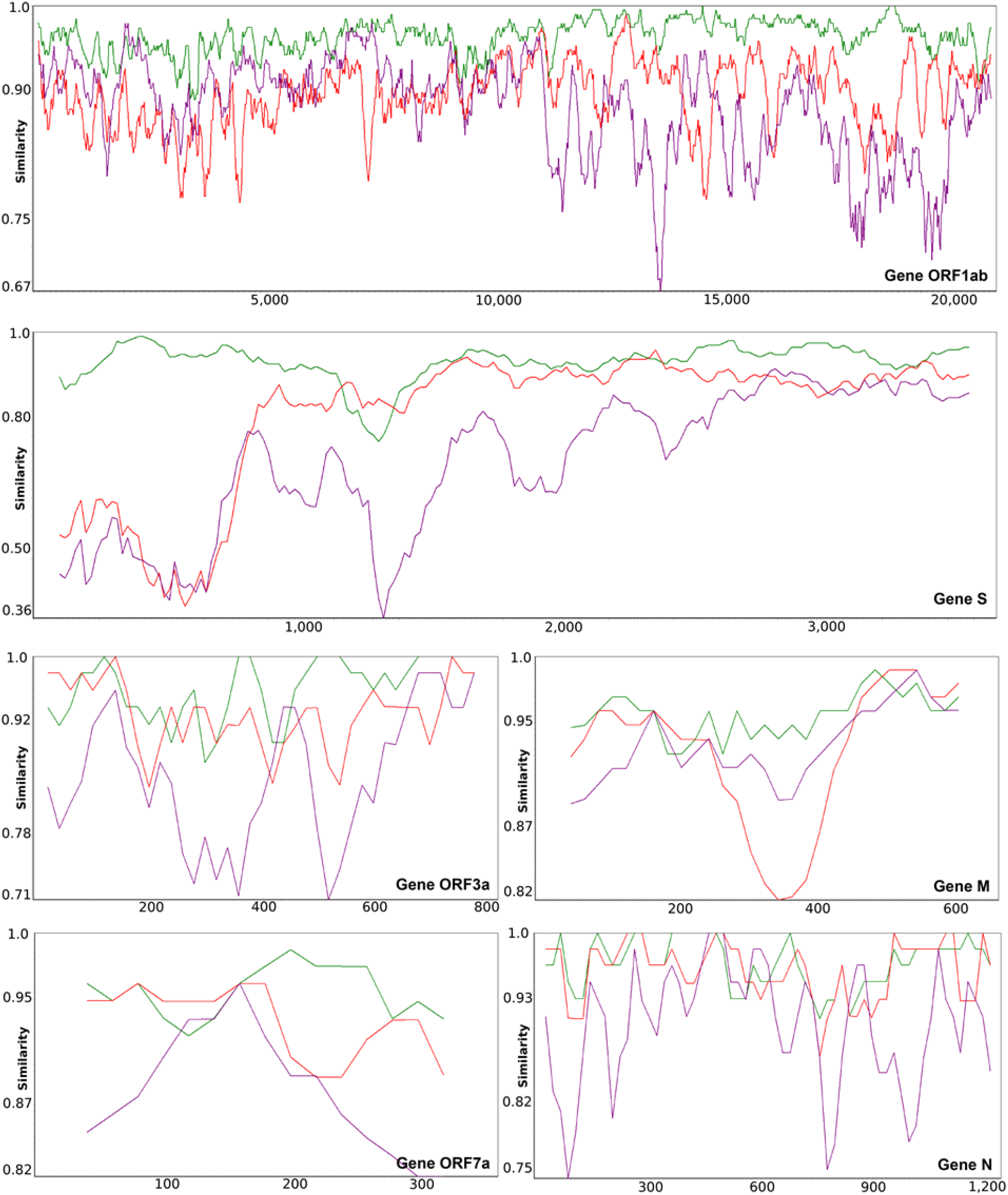
Gene-by-gene SimPlot similarity analysis performed to compare gene sequences of the Wuhan SARS-CoV-2 2020 reference genome with those of the RatTG13 genome (green), GD Pangolin CoV consensus genome (red) and Bat CoVZ consensus genome (violet). Graphs are shown for genes ORF1ab, S, ORF3a, M, ORF7a and N that encompass the most important overlaps between the RaTG13, GD Pangolin CoV and Bat CoVZ similarity curves.

### Φ-test recombination analysis

We conducted gene-by-gene Φ-test recombination analysis (Bruen et al. 2006) to further investigate eventual recent recombination events which may have occurred between orthologous gene sequences of the Wuhan SARS-CoV-2, RaTG13, GD Pangolin 1 CoV, GD Pangolin P2S CoV, bat CoV ZC45 and bat CoV ZXC21 viruses. The Φ-test was carried out with different sliding window sizes, varying from 50 to 400 (with a step of 50), and the window progress step of 1 as the window size can affect the test outcome. The results presented in Table 1 are reported for the window size corresponding to the smallest *p*-value found for a given gene. The *p*-values lower than or equal to the 0.05 threshold were considered as significant. They indicate the presence of recombination in the gene under study.

**Table 1.** The results of the Φ recombination test carried out for gene and whole genome sequences of Wuhan SARS-CoV-2, RaTG13, GD Pangolin 1 CoV, GD Pangolin P2S CoV, bat CoV ZC45 and bat CoV ZXC21.

| Region | Φ-test result<br>(p-value) | Recombination<br>detected<br>(yes/no) | Window<br>size |
| --- | --- | --- | --- |
| Gene ORF1ab | $2.23 \times 10^{-3}$ | Yes | 150 |
| RB domain | too short | - |  |
| Gene S | $8.60 \times 10^{-3}$ | Yes | 200 |
| Gene ORF3a | $1.13 \times 10^{-5}$ | Yes | 100 |
| Gene E | 0.56 | No | 200 |
| Gene M | 0.226 | No | 100 |
| Gene ORF6 | too short | - |  |
| Gene ORF7a | 0.00155 | Yes | 200 |
| Gene ORF7b | too short | - |  |
| Gene ORF8 | 0.0453 | Yes | 200 |
| Gene N | 0.024 | Yes | 50 |
| Gene ORF10 | too short | - |  |
| Whole genomes | 0.0156 | Yes | 200 |

According to the Φ-test (see Table 1), statistically significant recombination events involving these six coronaviruses have been detected in genes ORF1ab, S, ORF3a, ORF7a, ORF8, N and in the whole genome sequences. It is worth noting that recombination events in genes ORF1ab, S and ORF3a were detected with high confidence, with *p*-values of 0.0023, 0.0086 and 0.0000113, respectively.

### Horizontal gene transfer and recombination analyses

In addition to the SimPlot similarity and the Φ-test recombination analyses, we also inferred the whole genome phylogeny (Fig. 1b) and all individual gene trees for the main 11 CoV genes (Figs. 3a-k) as well as for the RB domain of the spike protein (Fig. 3l and Fig. 4a) using the RAxML method (Stamatakis 2006) with bootstrapping, and then conducted a detailed horizontal gene transfer-recombination analysis (Fig. 3) using the HGT-Detection program available on the T-Rex web server (Boc et al. 2012). It is worth noting that the presentation of the whole genome tree could be a bit confusing in our context, as the central premise of this work is that the evolution of coronavirus organisms is driven by the gene transfer and recombination mechanisms, but it remains necessary and serves as a support tree topology to represent the available species groups (Fig. 1b) as well as the detected gene transfer and recombination events (Figs. 3 and 4). The HGT-Detection program allows one to infer all possible horizontal gene transfer events for a given group of species by reconciling the species tree (i.e. whole genome tree in our case) with different gene phylogenies built for whole individual genes or some of their regions (Denamur et al. 2000; Boc and Makarenkov 2011). The bootstrap support of the inferred gene transfers was also assessed by HGT-Detection. It is worth noting that the bootstrap support of horizontal gene transfer events (HGT) is usually lower than that of the related branches of the species and gene phylogenies. For example, in order to get an HGT bootstrap support of 100%, gene transfer from cluster C1 to cluster C2 in the species tree must be present in all gene transfer scenarios inferred from all replicated multiple sequence alignments (MSAs) used in bootstrapping, and clusters C1 and C2 must be neighbor clusters in all gene trees inferred from replicated MSAs (Boc et al. 2010).

**Figure 3.**
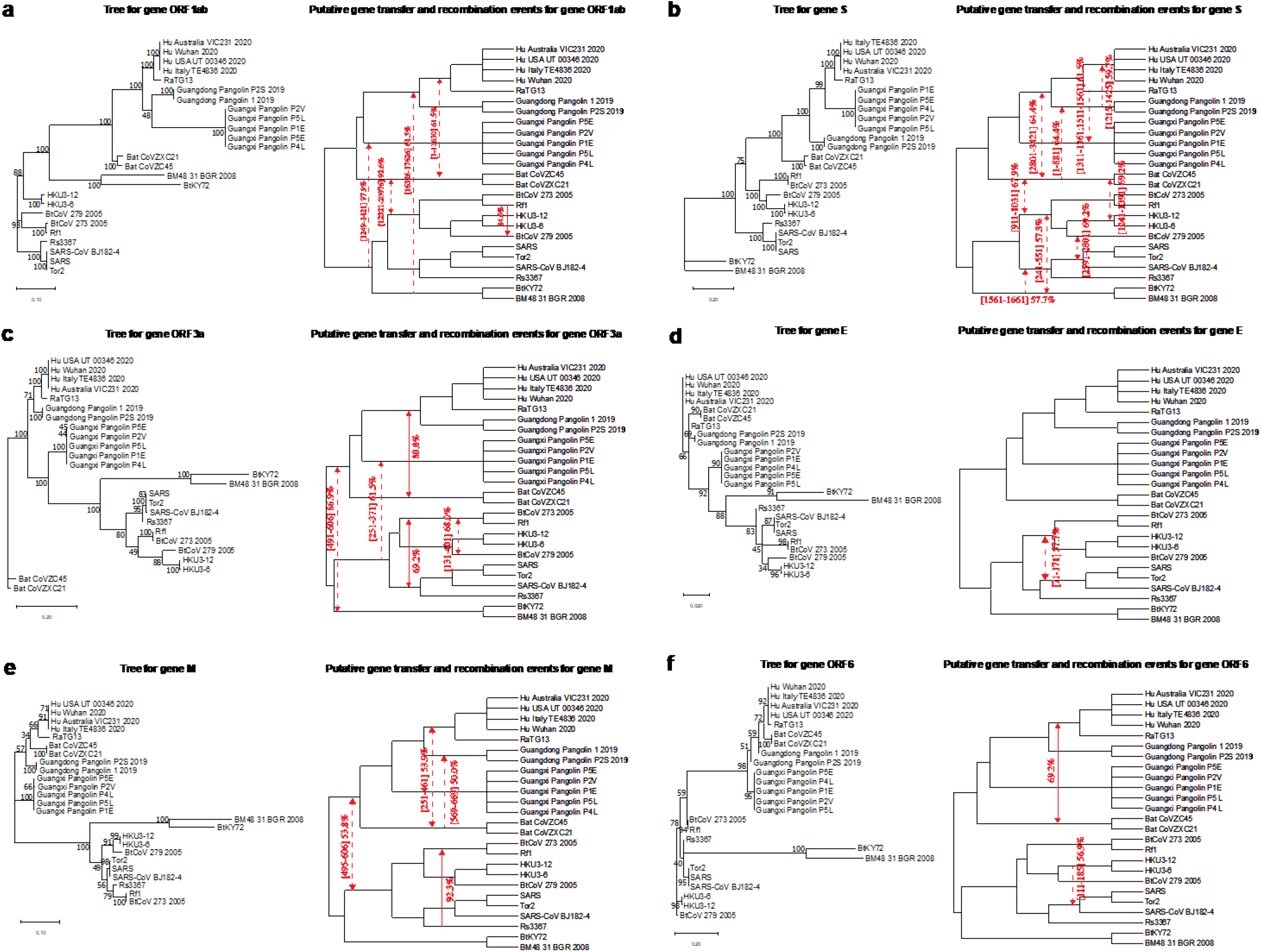
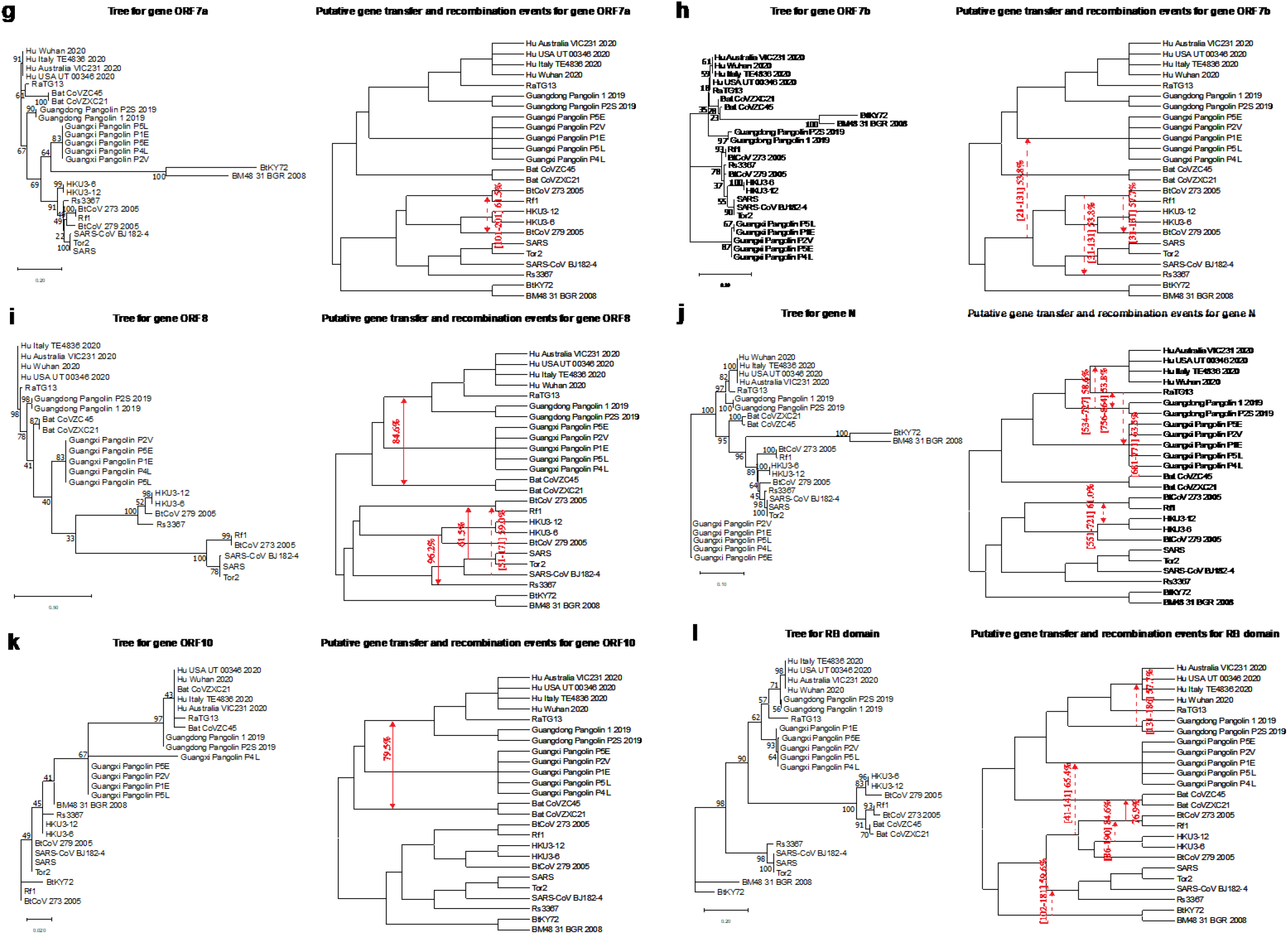
Putative horizontal gene transfer events found for the main 11 genes and the RB domain of 25 SARS-CoV and SARS-CoV-2-related viruses. Left part of each portion of the figure shows the gene tree, while its right part shows the species tree (i.e. whole genome tree) into which statistically significant horizontal gene transfers are mapped. Full lines represent complete gene transfers (i.e. when a complete copy of the donor’s gene is incorporated into the recipient genome; this accounts for intergenic recombination) and dashed lines represent partial gene transfers (i.e. when a mosaic gene is formed by recombination of two homologous gene sequences; this accounts for intragenic recombination). Numbers on the internal branches represent their bootstrap scores. Transfer directions are represented by arrows (when the direction is not certain, the arrow is bidirectional). Gene fragments transferred from the donor organism are indicated between brackets and followed by the transfer bootstrap score calculated by the Partial HGT-Detection program (Boc and Makarenkov, 2011).

**Figure 4.**
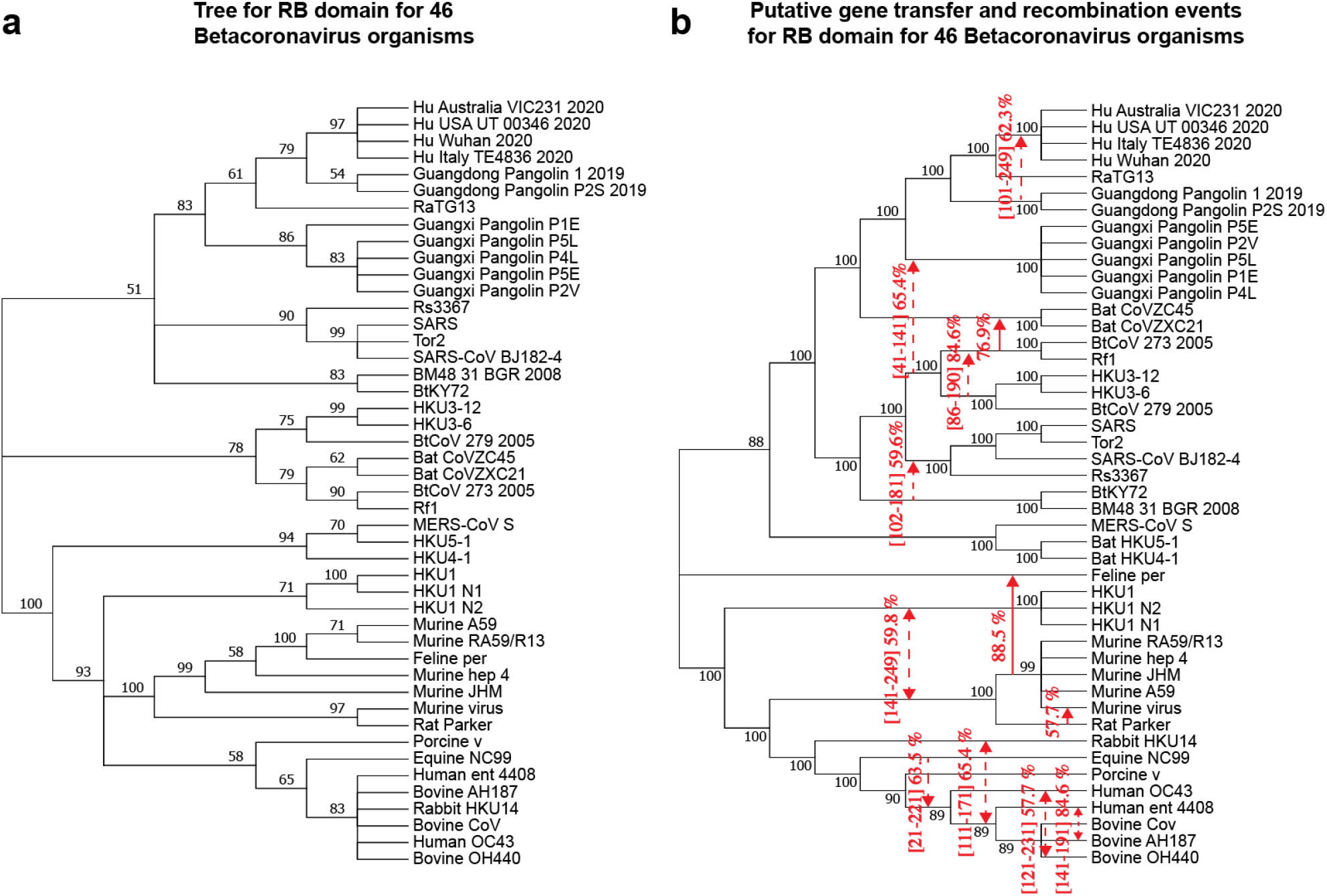
Putative horizontal gene transfer events found for the RB domain (amino acid sequences) of 46 betacoronavirus organisms (extended version of Fig. 3i). Left part of the figure presents the gene tree of the RB domain. Right part of the figure presents the species tree (i.e. whole genome tree) with putative horizontal gene transfers mapped into it. The explanations of the caption of Fig. 3 also apply for this figure.

Importantly, each detected horizontal gene transfer event can be interpreted in three ways: (1) It can represent a unique complete or partial HGT event involving distant species, as discussed above; (2) it can represent the phenomenon of parallel evolution, which the involved species might have undergone; and (3) it can also represent the situation where a new species (i.e. a gene transfer recipient) was created by recombination of the donor species genome with the genome of a recipient neighbor in the species phylogeny (this was potentially the case of the gene transfers from GD Pangolin CoV to SARS-CoV-2, which is a neighbor of RaTG13 in the species phylogeny, found for genes S and N; see our gene-by-gene analysis below).

We will now discuss the evolution of the main 11 CoV genes as well as that of the RB domain of the spike protein with emphasis on the specific horizontal gene transfer and recombination events detected for each of them.

### Evolution of gene ORF1ab

Gene ORF1ab occupies more than two thirds of coronavirus genomes. It encodes the replicase polyprotein, being translated from ORF1a (11826 to 13425 nt) and ORF1b (7983 to 8157 nt) (Woo et al. 2010), and plays an important role in virus pathogenesis (Graham et al. 2008).

The bootstrap support of the branches of the ORF1ab gene phylogeny is commonly very high, except for the branch connecting the clusters of GD and GX Pangolin CoVs (Fig. 3a). For this gene, we found a horizontal gene transfer-recombination event between the ancestors of the clusters of the Bat CoVZ organisms and the cluster including the RaTG13 and SARS-CoV-2 viruses. This event concerns a half of this long gene, occurring in gene region [1-11630] with bootstrap support of 61.5% (as this transfer does not affect the whole gene sequence, it is called *a partial HGT*). A transfer of the whole ORF1ab gene (i.e. *a complete HGT*) from the bat RF1 CoV to bat BtCoV 279 2005 has been detected with bootstrap score of 84.6%. The transfer between the cluster of the bat BTKY72 and BM48 31 BGR 2008 coronaviruses and the cluster involving SARS-CoV-2, RaTG13 and GD Pangolin CoVs has been detected on gene region [16326-17626] with bootstrap support of 61.5%. Finally, a deep phylogeny transfer from CoV organisms from the bottom part of the tree (i.e. SARS-like 2003-2013 viruses) to the cluster of GX Pangolins CoVs has been found in a relatively short region [1249-1421] with a high bootstrap score of 97.9%.

### Evolution of gene S

Gene S is among the most important coronavirus genes since it regulates the ability of the virus to overcome species barriers and allows interspecies transmission from animals to humans (Zhang et al. 2006). S proteins are responsible for the “spikes” present on the surface of coronaviruses, giving this virus family a specific crown-like appearance. These proteins are type I membrane glycoproteins with signal peptides used for receptor binding (Woo et al. 2010). They play a crucial role in viral attachment, fusion and entry, being a target for development of antibodies, entry inhibitors and vaccines (Tai et al. 2020).

As shown on the SimPlot diagram (Fig. 1a and Fig. 2), this gene has the most variable sequences among all genes of coronavirus genomes. The bootstrap scores of internal branches of the phylogeny of gene S are all very high (Fig. 3b). For this gene, a partial gene transfer was found in region [1-881] between the Bat CoVZ group and the cluster of GD Pangolin CoVs with bootstrap support of 64.4%. Another partial transfer from GD Pangolin CoVs to SARS-CoV-2 was found on the interval [1215-1425] with bootstrap score of 59.7%. This transfer corresponds to the RB domain. Partial transfers were detected from RaTG13 to GX Pangolin CoVs in regions [1311-1361] and [1511-1561] with an average bootstrap score of 61.5%. Finally, another partial transfer was found between the bat CoVZ group and the cluster including BtCoV 273 2005, Rf1, HKU3-12, HKU3-6 and BtCoV 279 2005.

### Evolution of gene ORF3a

The protein 3a is unique to SARS-CoV and SARS-CoV-2. It is essential for disease pathogenesis. In SARS-related CoVs, it forms a transmembrane homotetramer complex with ion channel function and modulates virus release (Lu et al. 2006). The sequences of ORF3a are as highly variable, almost as those of gene S. The bootstrap scores of the ORF3a gene tree (Fig. 3c) are usually high, except for that of the branch connecting the cluster of GD Pangolin CoVs and the cluster of RatTG13 and SARS-CoV-2 (71%), and the branch connecting clusters of some old CoVs at the bottom part of the tree.

For this gene, a full gene transfer was found between the cluster of SARS-CoV-2, RaTG13 and GD Pangolin CoVs, and the group of Bat CoVZ viruses, with statistical significance of 80.8%. A partial gene transfer was detected between the lower part of the tree (i.e. SARS-like 2003-2013 viruses), excluding bat viruses from Kenya and Bulgaria (i.e. BtKY72 and BM48 31 BGR 2008), and the GX Pangolin CoVs in region [251-371] with bootstrap support of 61.5%. The remaining identified transfers were a deep phylogeny transfer and two transfers between CoVs from the previous SARS outbreak.

### Evolution of gene E

This gene is among the shortest in the SARS-CoV-2 genome. The protein E is a small transmembrane protein associated with the envelope of coronaviruses. It is well conserved among all CoVs (Fig. 3d). Gene E is usually not a good target for phylogenetic analysis because of its short sequence length (Woo et al. 2010). The bootstrap scores of its gene phylogeny (Fig. 3d) are mostly mediocre. A single gene transfer-recombination event has been found for this gene. It affects region [71-171] and involves CoVs from the previous SARS outbreak.

### Evolution of gene M

This gene is particularly important since it is responsible for assembly of new virus particles (Kandeel et al. 2020). Bootstrap scores of the gene tree are on average much lower than those of genes ORF1ab, S and ORF3a (Fig. 3e).

Here, we detected two partial gene transfers involving the viruses of the Bat CoVZ group and affecting: (1) the SARS-CoV-2 and RaTG13 CoVs in region [251-461] with bootstrap support of 53.9%, and (2) the GD Pangolin CoVs in region [569-669] with bootstrap support of 50%. Moreover, a complete gene transfer from Rs367 to the cluster of Rf1 and BtCoV 273 2005 was found for this gene with bootstrap score of 92.3%.

### Evolution of gene ORF6

Gene ORF6 impacts the expression of transgenes (Mortiboys et al. 2015). The gene phylogeny of ORF6 (Fig. 3f) has multiple internal branches with low bootstrap support. For this gene, we found a complete gene transfer between the Bat CoVZ group and the cluster including SARS-CoV-2 and RaTG13 with bootstrap support of 69.2%. Furthermore, a partial gene transfer between HKU3-12 CoV and the SARS-CoV cluster in region [111-185] with bootstrap score of 67.7% was also found for this gene.

### Evolution of genes ORF7a and ORF7b

The proteins encoded by coronavirus genes ORF7a and 7b have been demonstrated to have proapoptotic activity when expressed from cDNA (Schaecher et al. 2007). The phylogenies of these short genes are very unresolved (Figs. 3g and 3h). A gene transfer-recombination event found for ORF7b involves CoVs from the lower part of the tree, excluding CoVs of Kenyan and Bulgarian bats, and the GX Pangolin CoVs. It occurred in region [21-131] with bootstrap score of 53.8%. Interestingly, a transfer affecting the bat CoVs related to the previous SARS outbreak was found in both of these genes.

### Evolution of gene ORF8

It has been recently shown that the SARS-CoV-2 viral protein encoded from gene ORF8 shares the least homology with SARS-CoV among all the viral proteins, and that it can directly interact with MHC-I molecules, significantly downregulating their surface expression on various cell types (Zhang et al. 2020). The gene phylogeny of ORF8 has several internal branches with low bootstrap scores (Fig. 3i). It has been established that ORF8 protein of SARS-CoV has been acquired through recombination from SARS-related coronaviruses from greater horseshoe bats (Lau et al. 2015). A complete gene transfer was found for this gene between the cluster of SARS-CoV-2, RaTG13 and GX Pangolin CoVs and the bat CoVZ group with bootstrap support of 84.6%. The other detected transfers concerned the viruses of the SARS-CoV group.

### Evolution of gene N

The nucleocapsid protein (N) is one of the most important structural components of SARS-related coronaviruses. The primary function of this protein is to encapsulate the viral genome. It is involved in the formation of the ribonucleoprotein through interaction with the viral RNA (Kandeel et al. 2020). Several studies report that the protein N interferes with different cellular pathways, thus being a crucial regulatory component of the virus as well (Surjit et al. 2008). The phylogeny of gene N (Fig. 3j) is usually well resolved, except for two clusters in the SARS-CoV part of the tree with bootstrap scores of 64% and 45%. Our SimPlot analysis showed that the SARS-CoV-2 gene sequence of gene N is almost as similar to the RatTG13 gene sequence as it is to the GD Pangolin gene sequence. Precisely, for gene N, the Wuhan SARS-CoV-2 and RatTG13 viruses share 96.9% of the whole-gene identity, while the Wuhan SARS-CoV-2 and GD Pangolin CoV gene sequences are 96.19% identical.

Three statistically significant gene transfer-recombination events have been detected for this gene. The most interesting of them is the partial gene transfer from the cluster of GD Pangolin CoVs towards the cluster of SARS-CoV-2 found in region [534-727] with bootstrap support of 58.6%. Another partial transfer, between RaTG13 and GD Pangolin CoVs, was detected in region [756-864] with bootstrap support of 53.8%. Finally, a partial transfer between the cluster containing the BtCoV 273 2005 and Rf1 viruses and the cluster of HKU CoVs was found in region [551-721] with bootstrap score of 61.0%.

### Evolution of gene ORF10

The protein ORF10 of SARS-CoV-2 includes 38-amino acids and its function is unknown (Yoshimoto 2020). The phylogeny of this short gene is not well resolved (Fig. 3k) with several internal branches having bootstrap support under 50%. The only complete gene transfer-recombination event detected for this gene affects the Bat CoVZ virus group and the cluster of SARS-CoV-2, RatTG13 and GD Pangolin CoVs. Its bootstrap score is 79.5%.

### Evolution of the RB domain

We carried out an independent analysis of the main evolutionary events characterizing the RB domain of the spike protein because of its major evolutionary importance. The S protein mediates the entry of the virus into the cells of the host species by binding to a host receptor through the RB domain located in its S1 subunit, and then merging the viral and host membranes in the S2 subunit. As SARS-CoV, SARS-CoV-2 also recognizes ACE2 as its host receptor binding to the S protein of the virus. Thus, the RB domain of the SARS-CoV-2 S protein is the most important target for the development of virus attachment inhibitors and vaccines (Tai et al. 2020).

We first studied the evolution of the RB domain for the 25 original organisms that are strongly phylogenetically related to SARS-CoV and SARS-CoV-2. The RB domain amino acid phylogeny (Fig. 3l) commonly exhibits high bootstrap scores of its internal branches, except for the root branch of the cluster of GD Pangolin CoVs (56%), the branch separating RatTG13 from the cluster of SARS-CoV-2 and GD Pangolin CoVs (57%), and the branch separating the cluster of GX Pangolin CoVs from the cluster including the SARS-CoV-2, GD Pangolin CoV and RatTG13 viruses (62%). Thus, the location of both pangolin clusters and that of RatTG13 are the most uncertain in this tree.

Our similarity analysis showed that SARS-CoV-2 and RatTG13 share 89.47% of the whole-gene identity, while the SARS-CoV-2 and GD Pangolin CoV RB domain amino acid sequences are 97.36% identical. Consequently, some exchange of genetic material between the clusters of GD Pangolin CoVs and SARS-CoV-2 would be expected here. In fact, a gene transfer from GD Pangolin CoVs to SARS-CoV-2 was detected in region [131-186] with bootstrap support of 57.7%. Another transfer found for this tree was that from the cluster containing BtCoV 273 2005, Rf1, HKU3-12, HKU3-6 and BtCoV 279 2005 to GX Pangolin CoVs in region [41-141] with bootstrap support of 65.4%.

Moreover, we also inferred an extended version of the RB domain tree, adding to it 21 coronavirus organisms (Fig. 4a) labeled as common cold CoV in the Gisaid coronavirus tree (Shu and McCauley 2017) and other coronavirus organisms available in GenBank (Prabakaran al. 2006). These additional CoVs include Human betacoronavirus 2c EMC/2012 (i.e. MERS-CoV S), Human coronavirus (i.e. HKU1 and its isolates N1 and N2), Human coronavirus OC43 and Human enteric coronavirus strain 4408. This extended analysis allowed us to discover some additional intergenic and intragenic gene transfer-recombination events involving human-related coronaviruses, including the exchange of genetic material between: (1) Murine CoVs and HKU1-related viruses, i.e. a complete gene transfer detected with high bootstrap score of 88.5%, and a partial transfer detected with bootstrap score of 59.8%; (2) Equine CoV and the cluster including Human OC43 and Enteric CoVs, i.e. a partial gene transfer with bootstrap score of 63.5%; (3) Rabbit HKU14 CoV and the cluster including the Enteric human CoV, and the three bovine CoVs, i.e. a partial gene transfer with bootstrap score of 65.4%; (4) Human OC43 CoV and Bovine OH440 CoV, i.e. a partial gene transfer with bootstrap score of 57.7%; and (5) Human Enteric CoV and Bovine AH187 CoV, i.e. a partial gene transfer with bootstrap score of 84.6%. Interestingly, no any gene transfer-recombination event between the viruses of the upper part of the species tree, containing SARS-related coronaviruses (Fig. 4b), and the lower part of this tree, containing MERS-related, HKU1-related, OC43 CoV-related and Enteric CoV-related viruses, has been detected.

### Analysis of intergenic recombination in 46-species phylogeny

We also carried out an analysis of intergenic (complete gene transfers) recombination events in the 46-species phylogeny discussed in the previous section. This extended analysis allowed us to discover some additional gene transfer-recombination events that affected the lower part of this larger phylogeny (see Fig. 5), including the 25 SARS-CoV-2-related viruses from Fig. 1b as well as MERS-CoV, HKU1 coronavirus strains, Human coronavirus OC43, Human enteric coronavirus, and different Bat, Rat Parker, Murine, Feline, Equine, Bovine, Porcine and Rabbit coronavirus strains. The analysis was conducted using the HGT-Detection program (Boc et al. 2010) available on the T-Rex web server (Boc et al. 2010). The transfers of the higher part of the tree presented in Figure 5 are the complete gene transfers from Figure 3. The most significant transfers found for the lower part of the tree include those between: (1) Murine JHM and Rat Parker coronaviruses found for gene N (bootstrap score of 55.4%); (2) Feline CoV and the ancestor of the cluster comprising Porcine, Human OC43, Human enteric and three Bovine CoVs for gene N2 (bootstrap score of 72.3%); (3) Rabbit CoV and the ancestor of the cluster comprising Human OC43, Human enteric and three Bovine CoVs for gene S (bootstrap score of 58.4%); and, finally, (4) an interesting complete transfer affecting the cluster of MERS-CoV S and two of its close relatives, i.e. bat coronaviruses HKU-4 and HKU-5, for gene ORF3a (bootstrap score of 76.4%). Gene transfer accounting for this recombination event most likely stems from an external source (a CoV organism which is absent in the tree). It is marked by a green arrow in Figure 5. It is worth noting that no intergenic recombination events between coronaviruses from the higher and lower parts of the extended CoV species phylogeny tree were found.

**Figure 5.**
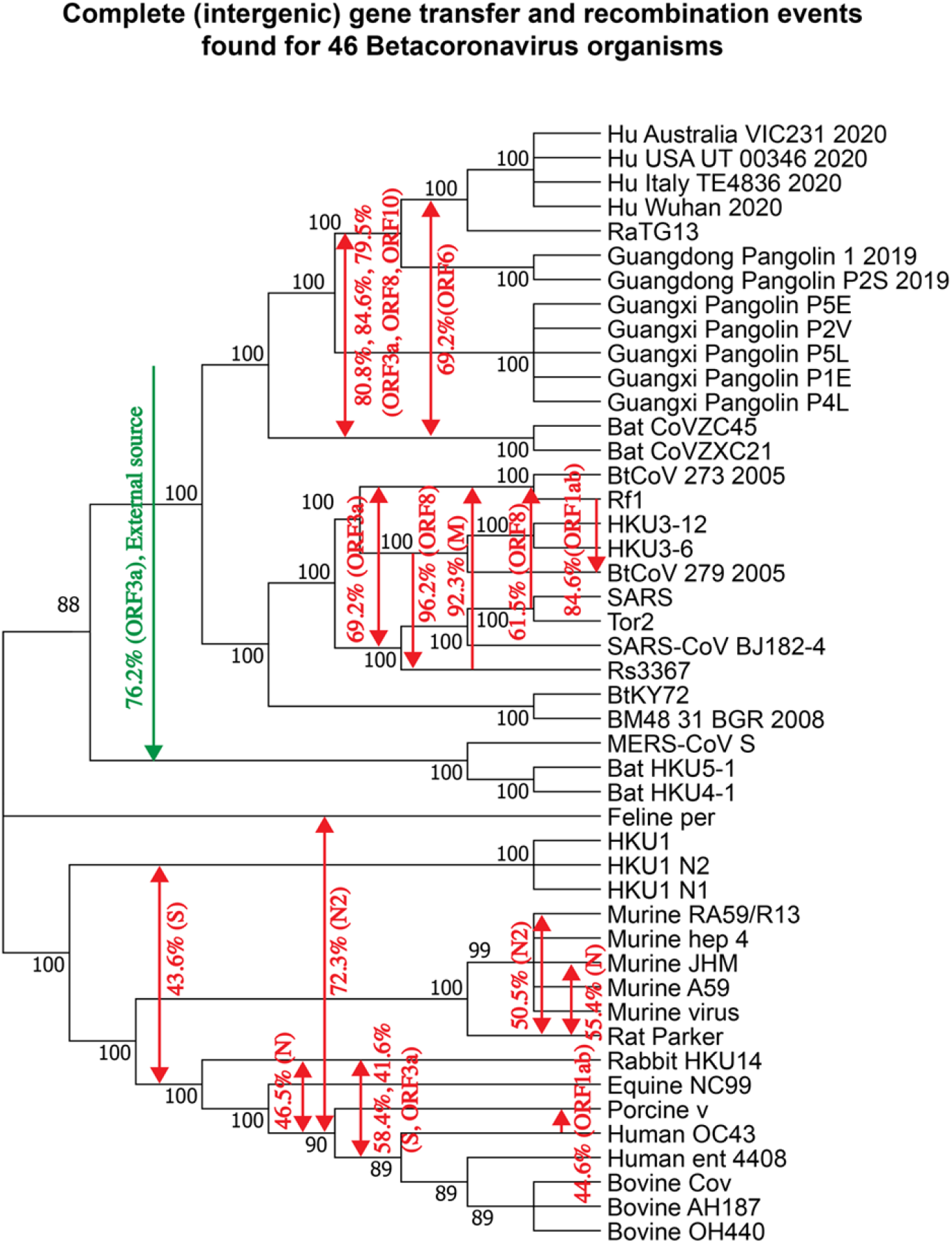
Putative complete horizontal gene transfer events (accounting for intergenic recombination) found for the main 11 genes of 46 betacoronavirus organisms from Fig. 4. The detected complete horizontal gene transfers are mapped into the species tree (i.e. whole genome tree). Numbers on the internal tree branches represent their bootstrap scores. Transfer directions are represented by arrows (when the direction is not certain, the arrow is bidirectional). Transfer bootstrap scores calculated by the HGT-Detection program (Boc et al. 2010) and the associated gene names are indicated on arrows.

### Cluster analysis of CoV gene phylogenies

Finally, we also carried out the CoV gene tree clustering to identify genes following similar evolutionary patterns (i.e. similar gene tree topologies). This analysis was performed using the k-means-based tree clustering algorithm adapted for clustering phylogenies with different numbers of leaves (Tahiri et al. 2018) as some gene trees contained less than 25 species (see the Methods section). Our results indicate that coronavirus genes followed three different patterns of evolution as the phylogenies of 11 CoV genes and that of the RB domain of the spike protein were partitioned into 3 disjoint clusters. Figure 6 presents the consensus trees of the detected tree clusters. The first of them (Fig. 6a) obtained using the *Consense* program from the Phylip package (Felsenstein 1993) is the extended majority rule consensus tree of the gene phylogenies of ORF1ab, S, RB domain of S and N. The supertree inferred by CLANN (Creeve and McInerney 2005) (in this case we could not use *Consense* since the species BM48_31_BGR_2008 and BtKY72 were missing in the gene phylogeny of ORF8) for the gene phylogenies of ORF3a, E, M, ORF6, ORF7a, ORF7b and ORF8 is shown in Fig. 6b. The consensus tree of the third cluster, containing a unique representative - gene tree of ORF10, is its RAxML gene phylogeny (Fig. 6c). attract

**Figure 6.**
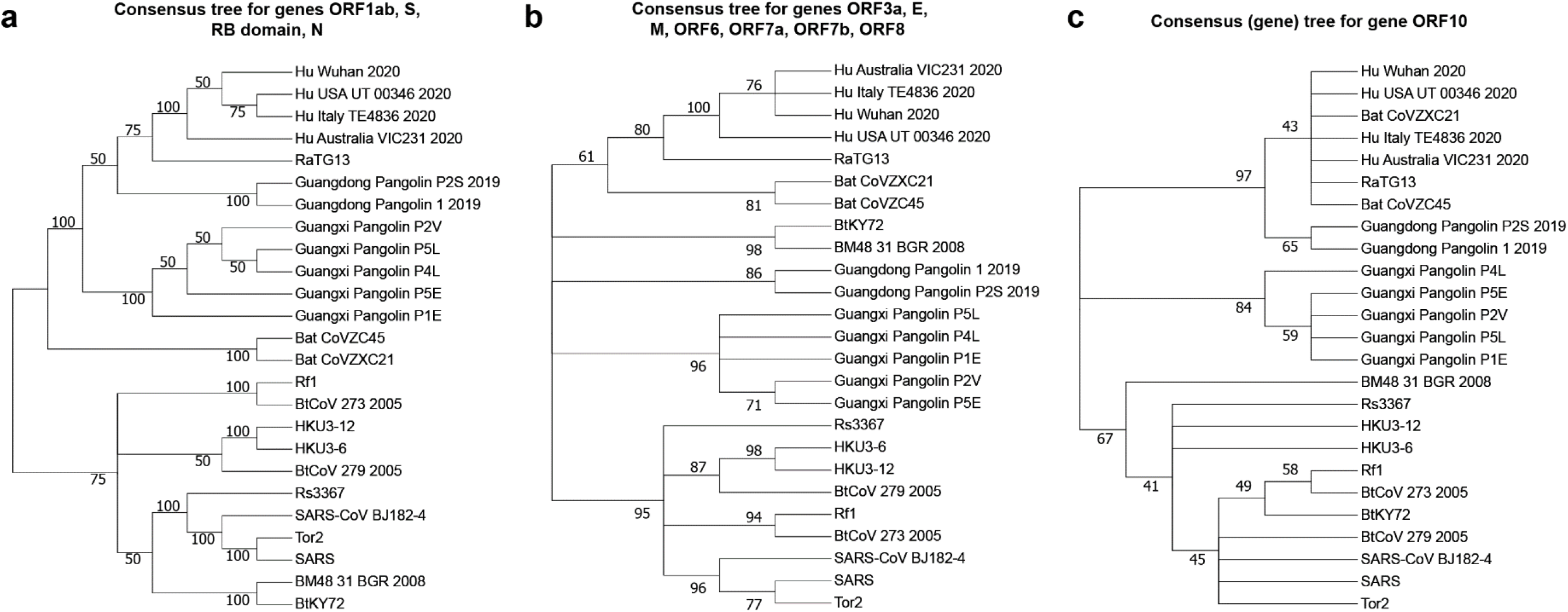
Three consensus trees showing the three ways of evolution of the SARS-CoV and SARS-CoV-2-related genes. Tree clustering was carried out using the k-means-based tree partitioning algorithm adapted for clustering trees with different numbers of leaves. The first tree is the extended majority-rule consensus tree inferred for the phylogenies of genes ORF1ab, S, RB domain of S and N, forming Cluster 1. This consensus tree was obtained using the *Consense* program from the Phylip package. The second tree is the best heuristic search (*hs*) CLANN supertree inferred for the phylogenies of genes ORF3a, E, M, ORF6, ORF7a, ORF7b and ORF8 (these gene phylogenies had different numbers of leaves), forming Cluster 2. The consensus tree of gene ORF10 is its RAxML tree, which was unique in Cluster 3. Bootstrap scores are indicated on the internal tree branches. Branches with bootstrap support lower than 40% were collapsed.

## Discussion

Recombination is a prevalent process contributing to diversity of most viruses, including SARS-CoV-2 and other betacoronavirus organisms. It allows viruses to overcome selective pressure and adapt to new hosts and environments (Pérez-Losada et al. 2015). In this work, we conducted thorough gene-by-gene horizontal gene transfer and recombination analysis of SARS-CoV-2-related viruses. Even though gene borders are not always a natural demarcation of regions where recombination might occur, e.g. two large portions of gene ORF1ab exhibit different evolutionary histories, the performed sliding window analysis allowed us to treat each considered genetic segment as an independent gene region having its own evolutionary history. We first performed a comparative analysis of four strains of SARS-CoV-2 with 21 close members of the SARS-CoV family, which revealed multiple horizontal gene transfer and recombination events between these coronavirus organisms. The most striking of them were statistically significant gene transfers from Guangdong Pangolin CoV to SARS-CoV-2 found in gene S (i.e. this transfer most likely accounts for a putative recombination event between Guangdong Pangolin CoV and RatTG13 in the RB domain region of the spike protein) and gene N (i.e. this transfer most likely accounts for a putative recombination event between Guangdong Pangolin CoV and RatTG13 in region [534-727] of this gene). These findings are in support of the hypothesis that SARS-CoV-2 genome is a chimera resulting from recombination of the RatTG13 and Guangdong Pangolin CoV genomes. According to some recent studies, the discovery of SARS-CoV-like coronaviruses from pangolins with similar RB domain, provides the most parsimonious explanation of how SARS-CoV-2 could acquire it via recombination or mutation (Anderson et al. 2020; Xiao et al. 2020; Li et al. 2020). The fact that the highlighted gene transfer-recombination events were detected only between Guangdong Pangolin CoV and SARS-CoV-2, but not between Guangxi Pangolin CoV and SARS-CoV-2, is another argument in favor of a mosaic recombinant origin of the SARS-CoV-2 genome, and against the parallel evolution paradigm. The confirmed recombination events in genes S and N (Fig. 3b and j) as well as the SimPlot recombination analysis of genes S, ORF3a, M, ORF 7a and N (Fig. 2) suggest that Guangdong pangolin is a likely intermediate host of SARS-CoV-2, on its way of transmission from bats to humans. The bat RatTG13 virus strain could infect a Guangdong pangolin, which was already a bearer of its own CoV virus, probably similar to that found in Guangxi pangolins. The two CoV genomes could then recombine and the resulting recombinant evolve into a SARS-CoV-2 mosaic strain which was transmitted to humans. Furthermore, we also discovered multiple gene exchanges between the cluster including the bat CoVZC45 and CoVZXC21 viruses, and the cluster including the SARS-CoV-2 strains and RatTG13. Some statistically significant gene transfer-recombination events between these CoV clusters were found in 6 of 11 coronavirus genes, namely in ORF1ab, ORF3a, M, ORF6, ORF8 and ORF10 (according to the gene transfer-recombination analysis conducted with HGT-Detection; see Fig. 3), and in genes ORF1ab, S, ORF3a, ORF7a, ORF8 and N (according to recombination analysis conducted with Φ-test; see Table 1). These findings confirm that not only the RatTG13 and GD Pangolin CoVs have influenced the evolution of SARS-CoV-2, but also the bat CoVZC45 and CoVZXC21 coronaviruses or their common ancestor.

The intergenic recombination analysis of the extended 46-species coronavirus phylogeny allowed us to detect eight additional statistically significant gene transfer events affecting the lower part of the tree. Among them, we need to highlight a complete gene transfer affecting the cluster of MERS-CoV S and two bat coronaviruses (HKU-4 and HKU-5), stemming from an external source, which was found for gene ORF3a (see Fig. 5). No recombination events between coronaviruses from the higher and lower parts of the extended CoV phylogeny were found (see Figs. 4b and 5). This finding suggests that the gene transfer-recombination history of coronaviruses from the SARS-Cov-2 cluster (common cold CoVs from the higher part of the extended 46-species phylogeny) and that of coronaviruses from the cluster including MERS-related, HKU1-related, OC43-related and Enteric-related CoVs (lower part of the extended 46-species phylogeny) can be studied separately.

It is worth noting that we have also tried to add to our data set some extra SARS-CoV-2 genomes (in addition to the four originally considered SARS-CoV-2 genomes from Wuhan in China, Italy, Australia and USA), but realized that it did not lead to discovery of any further gene transfer-recombination event in which these extra SARS-CoV-2 organisms could be involved. This happens because of a very high sequence similarity between the available SARS-CoV-2 genomes. For example, the root genomes of the two main SARS-CoV-2 lineages, A and B (according to a recent SARS-CoV-2 sequence classification proposed by Rambaut et al. 2020), share 99.89% of whole genome identity (we measured it between the following coronavirus organisms: Lineage_A_EPI_ISL_406801 and Lineage_B_MN908947.3; see Rambaut et al. 2020 for more details), while the Hu_Wuhan_2020 and Hu_Australia_VIC231_2020 SARS-CoV-2 genomes analyzed in our work share 99.98% of whole genome identity. The difference between the existing SARS-Cov-2 genomes can be mainly explained by the presence of particular sets of mutations, with respect to the root sequence (Rambaut et al. 2020). These mutations are few in numbers and are usually not contiguous. They can be detected by a simple genome comparison, and do not necessitate the use of horizontal gene transfer and recombination detection methods. It should be mentioned that when extra SARS-CoV-2 genomes were added to the 46 betacoronavirus organisms analyzed in our work, they were always involved into the same gene transfer-recombination events as the four originally considered SARS-CoV-2 genomes.

We also conducted a cluster analysis of the coronavirus gene trees in order to identify genes with similar evolutionary histories. This analysis revealed the presence of three clusters of gene phylogenies: the first cluster includes the phylogenies of genes ORF1ab, S, RB domain and N, the second cluster includes the phylogenies of genes ORF3a, E, M, ORF6, ORF7a, ORF7b and ORF8, and the third cluster contains only the phylogeny of gene ORF10. For example, the phylogenies of genes S and N, whose evolution was affected by the highlighted recombination events between Guangdong Pangolin CoV and RatTG13, were assigned to the same cluster.

## Conclusions

The main finding of our work is a detailed list of statistically significant horizontal gene transfer and recombination events inferred for 11 main genes and the RB domain of the spike protein of SARS-Cov-2 and related betacoronavirus genomes (see Figs. 3, 4 and 5). The main advantages of the conducted gene transfer and recombination analysis, compared to other recent works in the field (Lam et al. 2020; Zhang et al. 2020; Boni et al. 2020), is that it allowed us not only to identify genes and genomes that have been affected by recombination, but also to determine the donor and recipient organisms for each detected recombination event, to find out whether this event was intergenic or intragenic, and to assess its statistical significance via a bootstrap score. Our detailed horizontal gene transfer and recombination analysis was conducted for each of 11 main genes of the SARS-Cov-2 genome, involving 46 betacoronavirus organisms. The obtained results (see Figs. 1 to 5) suggest that SARS-Cov-2 could not only be a chimera resulting from recombination of the bat RaTG13 and Guangdong pangolin coronaviruses but also a close relative of the bat CoV ZC45 and ZXC21 virus strains. They also indicate that a Guangdong pangolin may be an intermediate host of SARS-CoV-2 prior to its transmission to humans. Furthermore, our topology-based clustering analysis of coronavirus gene trees revealed a three-way evolution of SARS-CoV-2 genes (see Fig. 6). It is worth mentioning that some incongruencies among horizontal gene transfer and recombination detection methods may exist when applied to the same data (Becq et al. 2010). In this work, we used three different methods for detecting gene transfer and recombination events (Φ-test recombination analysis of Bruen et al. 2006, intergenic HGT analysis of Boc et al. 2010, and both intergenic and intragenic HGT analysis of Boc and Makarenkov 2011) to study the evolution SARS-CoV-2 and related betacoronaviruses. While the method of horizontal gene transfer analysis of Boc et al. (2010) is based on the comparison of the species and gene tree topologies, that of Boc and Makarenkov (2011) conducts a sliding window analysis of aligned gene sequences. Both of them make part of a phylogenetic approach. In the future, it would be interesting to validate our horizontal gene transfer and recombination results using composition-based ("parametric") horizontal gene transfer detection methods (Ravenhall et al. 2015).

## Methods

### Data description

We explored the evolution of 25 coronavirus organisms, including a cluster of four SARS-CoV-2 genomes (from Wuhan in China, Italy, Australia and USA taken from different clusters of the Gisaid human coronavirus tree available at: https://www.gisaid.org; Shu and McCauley 2017), two GD Pangolin CoV genomes (obtained from dead Malayan pangolins during an anti-smuggling operation in the Guangdong province of China) and five Guangxi Pangolin CoV genomes (obtained from the Beijing Institute of Microbiology and Epidemiology). We also included in our analysis the RaTG13 bat CoV genome from *Rhinolophus affinis* from the Yunnan province of China, along with the cluster of two Bat CoVZ organisms, comprising the bat CoV ZC45 and ZXC21 viruses, collected in the Zhejiang province of China in 2018 and five bat CoV genomes sampled in bats across multiple provinces of China from 2006 to 2010 (denoted as BtCoV 273 2005, Rf1, HKU3-12, HKU3-6 and BtCoV 279 2005 in the whole genome CoV phylogeny, see Fig. 1b). Finally, we also considered the SARS-CoV-related genomes from the first SARS outbreak (i.e. human SARS, Tor2, SARS-CoV BJ182-4 CoVs and bat Rs3367 CoV found in *Rhinolophus sinicus*) and two CoV strains coming from bats found in Kenya and Bulgaria (BtKY72 and BM48 31 BGR 2008). Most of these CoV genomes have been originally considered by Lam et al. (2020). Moreover, for an extended analysis of putative gene transfer-recombination events affecting the RB domain of the spike protein and intergenic recombination events (complete gene transfers) affecting all the genes under study, we considered 21 additional coronavirus organisms, including viruses labeled as common cold CoVs in the Gisaid coronavirus tree (Shu and McCauley, 2017) and other CoV organisms studied by Prabakaran et al. (2006; they are available in GenBank (Benson et al. 2012) at: https://www.ncbi.nlm.nih.gov/Structure/cdd/PF09408). Supplementary Table 1 in Supplementary Material reports full organism names, host species and GenBank or Gisaid accession numbers for all CoV genomes analyzed in this study. We first carried out the SimPlot (Lole et al. 1999) similarity analysis of coronaviruses most closely related to SARS-CoV-2, comparing the Wuhan SARS-CoV-2 reference genome to a consensus genomes of five CoV groups (GD Pangolin CoVs, GX Pangolin CoVs, RaTG13, Bat CoVZ and Bat SL-CoV; see Fig. 1a). The GD Pangolin group in our analysis consisted of two Guangdong Pangolin CoVs available in Gisaid (GD Pangolin 1 and GD Pangolin P2S in Fig. 1b). This explains differences in the results of our SimPlot similarity analysis with the results of Lam et al. (2020), who considered only the first of these GD Pangolin CoVs in their SimPlot analysis. To avoid possible inconsistency and program crashes during the SimPlot similarity analysis (Lole et al. 1999), Φ-test recombination analysis (Bruen et al. 2006) and horizontal gene transfer detection (Boc et al. 2011), we replaced missing nucleotides in the low-coverage regions of the GD Pangolin P2S CoV genome by the corresponding nucleotides of the GD Pangolin 1 CoV genome. This allowed us to better highlight intersections between the GD Pangolin CoV group similarity curve and the RaTG13 similarity curve (see Figs. 1a and Fig. 2), which may indicate the presence of gene transfer-recombination events between the GD Pangolin and RaTG13 coronaviruses. Multiple sequence alignments for all gene and genome CoV sequences (in the Fasta format) used in this study as well as all inferred phylogenetic trees (in the Newick format) are available at: http://www.info2.uqam.ca/~makarenkov_v/Supplementary_Material.zip.

### Methods details

The VGAS (Viral Genome Annotation System) tool (Zhang et al. 2019), designed to identify automatically viral genes and perform gene function annotation, was used to validate all CoV genes extracted from GenBank and Gisaid. Multiple sequence alignments for 11 CoV genes of the 25 original, and then 46 (for an extended analysis), betacoronavirus organisms (nucleotide sequences), and for the RB domain of the spike (S) protein (amino acids), were carried out using the MUSCLE algorithm (Edgar 2004) with default parameters of the MegaX package (version 10.1.7) (Kumar et al. 2018). These alignments were used to infer gene trees presented in Figs. 3 and 4 (left part of each portion of the figure). The whole genome CoV sequences for the original 25, and then 46 (for an extended analysis), betacoronavirus organisms were aligned in MegaX using the same version of MUSCLE. The whole genome alignments were used to infer species trees (see Figs. 1b and 4b). The alignment accuracy for all gene and genome alignments was verified manually base by base. The GBlocks tool (version 0.91b; Castresana 2000) from the Phylogeny.fr web server (Dereeper et al. 2008) was used to eliminate sites with large proportions of gaps. The less stringent correction option of GBlocks was used.

The maximum likelihood (ML) gene and genome phylogenies were inferred using the RAxML algorithm (version v0.9.0; Stamatakis 2006). Each tree was constructed under the best-fit DNA/amino acid substitution model found using MegaX, and available in RAxML, for the corresponding multiple sequence alignment. The best available substitution model for genes ORF1ab, S, N and the whole genomes was (GTR+G+I), for genes ORF3a, E, ORF6, ORF7a it was (HKY+G), for gene ORF7b it was (HKY+I), for gene ORF8 it was (HKY+G+I), for gene ORF10 it was (JC), and for the RB domain it was (WAG+G) (see Supplementary Table 2). In each case, the bootstrap scores of internal branches of all phylogenies were calculated using 100 bootstrap replicates. All gene and genome trees were originally drawn in MegaX.

The Partial HGT-Detection program (Boc and Makarenkov 2011) from the T-Rex web server (Boc et al. 2012) was used to infer directional horizontal gene transfer-recombination networks for 11 CoV genes and the RB domain of the spike protein (see Figs. 3 and 4b). Rooted ML genome tree of 25, and then 46, CoV organisms, playing the role of the species tree, and multiple sequence alignments (for 11 CoV genes and the RB domain of the spike protein) provided by MUSCLE, were used as input parameters of the Partial HGT-Detection and HGT-Detection programs (Boc et al. 2010; for the extended analysis of intergenic recombination events). The PhyML algorithm (Guindon et al. 2005) with 100 bootstrap replicates was carried out to infer trees from different gene regions for each position of the sliding window used in Partial HGT-Detection. This algorithm was carried out separately with sliding window sizes of 10, 25, 50 and 100 sites as well as with the whole sequence alignments (to infer gene transfers of whole genes). The sliding window advancement sizes (i.e. step sizes) of 1 (for short genes) and 10 (for long genes and whole genomes) sites was used in our analysis. Gene transfer-recombination events with bootstrap support of at least 50% identified by Partial HGT-Detection (see Figs. 3 - right portion of each panel, and 4b), and at least 40% identified by HGT-Detection (see Fig. 5), were represented by mapping them into the species tree. Some of these transfers may in fact be explained by the paradigm of parallel evolution when species sharing similar environment undergo similar mutations and develop similar traits. SimPlot v3.5. (Lole et al. 1999) was used to carry out a sliding window analysis and determine patterns of sequence similarity using as reference the Wuhan SARS-CoV-2 2020 genome. This genome was compared to the RatTG13 genome as well as to the consensus genomes of the Guangdong Pangolin CoV group, Guangxi Pangolin CoV group, Bat CoVZ group and Bat SL-VoC group (see Fig. 1a). These consensus genomes were the default consensus genomes generated by SimPlot v3.5 in order to represent a group of species. In addition, gene-by-gene SimPlot similarity analysis was performed to compare the genes of the Wuhan SARS-CoV-2 2020 reference genome with those of the RatTG13, Guangdong Pangolin CoV and Bat CoVZ group genomes (Fig. 2).

The Φ-recombination test (Bruen et al. 2006) was performed to detect recombination patterns among individual genes and whole genome sequences of the Wuhan SARS-CoV-2, RaTG13, GD Pangolin 1 CoV, GD Pangolin P2S CoV, CoV ZC45 and CoV ZXC21 viruses (see Table 1). The Φ-test was conducted with sliding windows of sizes 50 to 400 (with a step of 50) and the window progress step of 1. The version of the Φ-test used in our study was that provided by David Bryant at his web site: https://www.maths.otago.ac.nz/~dbryant/software.html.

Tree clustering (Fig. 6) was carried out using the k-means-based tree clustering algorithm adapted for clustering trees with different numbers of leaves (Tahiri et al. 2018) because some gene trees contained less than 25 species. The latest version of the tree clustering program was used (it is available at: https://github.com/TahiriNadia/KMeansSuperTreeClustering). The program was run with the following options - Tree clustering method: k-means; cluster validation index: Calinski-Harabasz; penalization parameter α = 0; Tree distance: Robinson and Foulds topological distance (not squared; Makarenkov and Leclerc 1996; Leclerc and Makarenkov 1998, Makarenkov and Leclerc 2000). The only difference with the default parameters of the program was that we set the penalization parameter α to 0 because 11 out 12 trees contained a full set of 25 species.

For the first cluster of trees (i.e. trees of genes ORF1ab, S, RB domain of S, and N) inferred for the full list of 25 species, the *Consense* program of the Phylip package (Felsenstein 1993) was used to infer the extended majority-rule consensus tree. As the sequences of BM48_31_BGR_2008 and BtKY72 were missing in the multiple sequence alignment of gene ORF8, we applied a supertree reconstruction method to retrace consensus evolutionary patterns for the cluster of genes ORF3a, E, M, ORF6, ORF7a, ORF7b, and ORF8. The *CLANN* program (Creevey and McInerney 2005) was used with the best heuristic search (*hs*) and *bootstrap* options with 100 replicates to infer a supertree for these genes. The consensus tree of the third cluster, containing the only tree of ORF10, is the ORF10 gene tree inferred with RAxML.

## Abbreviations

ACE2: Angiotensin-converting enzyme 2
CoV: Coronavirus
COVID-19: Coronavirus disease 2019
GD pangolin: Guangdong pangolin
GX pangolin: Guangxi pangolin
HGT: Horizontal gene transfer
MERS-CoV: Middle East respiratory syndrome coronavirus
MSA: multiple sequence alignment
ORF: Open reading frame
RB domain: Receptor binding domain
SARS: Severe acute respiratory syndrome
SARS-CoV: Severe acute respiratory syndrome coronavirus

## Declarations

### Ethics approval and consent to participate

All the experiments carried out in this study are in accordance with Canadian legislation, and the research performed does not require any ethical permits in Canada.

### Consent for publication

Not applicable.

### Availability of data and materials

The datasets supporting the conclusions of this article are available in our data archive at: http://www.info2.uqam.ca/~makarenkov_v/Supplementary_Material.zip and in Supplementary Material.

### Authors’ contributions

VM, GR and PL supervised and designed the study. BM and VM performed data processing, recombination and horizontal gene transfer analyses. All authors read and approved the final manuscript.

## Acknowledgements

We thank Compute Canada and Université du Québec à Montréal for providing us with necessary computational resources. We also thank Dr. Fernando Gonzalez-Candelas and two anonymous reviewers for their valuable comments on this manuscript.

